# ATP-gated P2x7 receptor is a major channel type at type II auditory nerves and required for hearing sensitivity efferent controlling and noise protection

**DOI:** 10.1101/2024.10.14.618333

**Authors:** Chun Liang, Tian-Ying Zhai, Shu Fang, Jin Chen, Li-Man Liu, Ning Yu, Hong-Bo Zhao

**Affiliations:** Department of Surgery – Otolaryngology, Yale University Medical School, 310 Cedar Street, New Haven, CT, USA, 06510; Department of Otolaryngology, University of Kentucky Medical Center, 800 Rose Street, Lexington, Kentucky, USA, 40536

**Author notes:** Corresponding Author: Hong-Bo Zhao, Ph.D./M.D., Professor, Dept. of Surgery – Otolaryngology, Yale University Medical School, 310 Cedar Street, New Haven, CT 06510, USA. These authors contributed equally to this work (Chun Liang, Tian-Ying Zhai, Shu Fang).

**Keywords:** type II spiral ganglion, LOC, MOC, cochlear efferent system, hearing oversensitivity, noise-induced hearing loss, outer hair cell electromotility, active cochlear mechanics

## Abstract

Hearing sensitivity and noise protection are mediated and determined by negative feedback of the cochlear efferent system. Type II auditory nerves (ANs) innervate outer hair cells (OHCs) in the cochlea and provide an input to this efferent control. However, little is known about underlying channel information. Here, we report that ATP-gated P2x7 receptor had a predominant expression at type II ANs and the synaptic areas under inner hair cells and OHCs with lateral and medial olivocochlear efferent nerves. Knockout (KO) of P2x7 increased hearing sensitivity with enhanced acoustic startle response (ASR), auditory brainstem response (ABR), and cochlear microphonics (CM) by increasing OHC electromotility, an active cochlear amplifier in mammals. P2x7 KO also increased susceptibility to noise. Middle level noise exposure could impair active cochlear mechanics resulting in permanent hearing loss in P2x7 KO mice. These data demonstrate that P2x7 receptors have a critical role in type II AN function and the cochlear efferent system to control hearing sensitivity; deficiency of P2x7 receptors can impair the cochlear efferent suppression leading to hearing oversensitivity and susceptibility to noise.

## Introduction

The descending cochlear efferent system provides a negative feedback loop to regulate hair cell activity, which plays an important role in the control of hearing sensitivity and protection from noise trauma (Guinan, 2006; Ciuman, 2010; Clause et al., 2017; Boero et al., 2018, Zhao et al., 2022). The cochlear efferent nerves are composed of the lateral olivocochlear (LOC) nerves and medial olivocochlear (MOC) nerves. The LOC nerves project from the lateral superior olivary nucleus in the brainstem to inner hair cells (IHCs) and form synapses with the dendrites of type I afferent auditory nerves (ANs) under the IHCs (Liberman et al., 1990; Maison et al., 2003). The MOC nerves originate from the medial superior olivary nucleus in the brainstem projecting to outer hair cells (OHCs) and supporting cells in the cochlea to mediate OHC electromotility (Nadol and Burgess, 1994; Zhao et al., 2022), which is an active cochlear amplifier to increase hearing sensitivity and frequency selectivity (Brownell et al., 1985; Zhao and Santos-Sacchi, 1999; Zheng et al., 2000; Liberman et al., 2002).

However, on the other hand, it is not so clear for input source(s) to the cochlear efferent system. It has been proposed that type II auditory nerves (ANs) may provide an important input to this negative feedback loop for controlling of OHC electromotility and eventually hearing sensitivity (Froud et al., 2015). Different from type I ANs, which innervate IHCs and occupy ∼95% of all ANs, Type II ANs innervate OHCs and only occupy ∼5% of ANs. It has been reported that type II ANs have important functions responsible for high-intensity sound stimulation (Weisz et al., 2009, 2014, 2021) and auditory nociception (Flores et al., 2015; Liu et al., 2015). However, since the type II AN is thin and rare and lacks specific cell markers, it is difficult to be identified and recorded. Currently, little is known about the underlying channel information.

ATP is an important extracellular signaling molecule and can activate purinergic (P2) receptors to influence physiological functions in many aspects (Jacobson et al., 2002; Surprenant and North, 2009). P2 receptors have two subgroups: ATP-gated ionotropic (P2x) receptors and G-protein coupled metabotropic (P2y) receptors. Each subgroup has several subtypes. In the cochlea, both P2x and P2y receptors have extensive expressions and play important roles in hearing in many aspects (Housley et al., 2009, Vlajkovic and Thorne, 2022). For example, P2x2 receptors have extensive expressions in the cochlea (Housley et al., 1999; Jarlebark et al., 2000). P2x2 mutations can induce hearing loss (Yan et al., 2013; Faletra et al., 2014; Zhu et al., 2017). The P2x7 receptor also expresses in the cochlea, particularly, in spiral ganglion (SG) neurons (Nikolic et al., 2002; Prades et al., 2021, Shi et al., 2024). However, its function remains unclear. In this study, we used transgenic mouse models and found that P2x7 has extensive expression at the type II SG neurons and the synapse areas of LOC and MOC nerves with IHCs and OHCs. Knockout of P2x7 led to deficiency of the negative cochlear efferent control and increased hair cell activity, therefore resulting in hearing over-sensitivity and vulnerability to noise. These data suggest that P2x7 receptors have a critical role in the functions of type II ANs and the cochlear efferent system.

## Materials and Methods

### Animals

P2x7 knockout (KO) mice were purchased from The Jackson Lab (Stock No: 005576) and were crossed with CBA/CaJ strain (Stock No: 000654, The Jackson Lab) for more than 4 generations. Both genders of adult mice were used in experiments. The *Peripherin-*eGFP mouse (Elliott et al., 2021) was obtained from Professor Ebenezer Yamoah (University of Nevada, USA). Mice were housed in a quiet individual room in basement with regular 12/12 light/dark cycle. The background noise level in the mouse room at mouse hearing range (4-70 kHz) was <30 dB SPL. All procedures and experiments followed in the use of animals were approved by the University of Kentucky’s Animal Care & Use Committee (UK: 00902M2005) and Yale University’s Institutional Animal Care & Use Committee (Yale: 2022-20463) and conformed to the standards of the NIH Guidelines for the Care and Use of Laboratory Animals.

### Immunofluorescent staining and confocal microscopy

As described in our previous reports (Zhao et al., 2021, 2022; Liu et al., 2023), the mouse cochlea was freshly isolated. The cochlea was incubated with 4% paraformaldehyde in PBS for 0.5-1 h. After fixation, the cochlea was dissected in the normal extracellular solution (NES) (142 NaCl, 5.37 KCl, 1.47 MgCl_2_, 2 CaCl_2_, 10 HEPES in mM, pH 7.4) or embedded in OCT for cross-section. The isolated cochlear sensory epithelia or the cochlear cross-sections were incubated in a blocking solution (10% goat serum and 1% BSA of PBS) with 0.1% Triton X-100 for 30 min. Then, the epithelia or sections were incubated with primary antibodies for P2x7 (1:250, rabbit anti-P2x7 polyclonal, #8232, Sigma-Aldrich, USA), Neurofilament (1:250, chicken anti-Neurofilament-H polyclonal, AB5539, Millipore), CtBP2 (1:500, mouse anti-CtBP2 IgG1, #612044, BD Bioscience), and eGFP (1:250, mouse anti-eGFP IgG2b, MA1-952, ThermoFisher) in the blocking solution at 4°C overnight. After washout, the epithelia were incubated with corresponding second Alexa Fluor antibody (1:400, goat anti-rabbit IgG Alexa Fluor 488, A-11034, goat anti-rabbit IgG Alexa Fluor 568, A-11036, goat anti-mouse IgG1 Alexa Fluor 568, A-21124, goat anti-chicken IgY Alexa Fluor 488, A-11039, goat anti-chicken IgY Alexa Fluor 568, A-11041, and goat anti-mouse IgG2b Alexa Fluor 488, A-21141, Thermo Fisher Sci) at room temperature (23 °C) for 2 hr. Before washout, 4’, 6-diamidino-2-phenylindole (DAPI, 50 μg/ml, D1306; Thermo Fisher Sci) was added to visualize the cell nuclei. After incubation for 2 min, the sensory epithelia were washed by PBS for 3 times and whole-mounted on the glass-slide. The stained epithelia or slides were observed under a Nikon A1R or AX confocal microscope system (Zhao et al., 2021, 2022).

### Acoustic startle response recording

Acoustic startle response (ASR) was recorded by use of a computer-controlled SR-LAB Startle Response System (San Diego Instruments, San Diego, CA). ASR was evoked by a series of white-noise pulses (25-ms duration) from 80 to 110 dB SPL in a 10 dB step.

### Auditory brainstem response and distortion product otoacoustic emission recording

Auditory brainstem response (ABR) was recorded by use of a Tucker-Davis R3 workstation with ES-1 high frequency speaker (Tucker-Davis Tech. Alachua, FL) (Zhu et al., 2013, 2015; Mei et al., 2017, Zhao et al., 2021, 2022; Liu et al., 2023). ABR was recorded by stimulation with clicks in alternative polarity and tone bursts (4-40 kHz) from 80 to 10 dB SPL in a 5 dB step in a double-walled sound isolation room. The signal was amplified (50,000x), filtered (300-3,000 Hz), and averaged by 500 times. The ABR threshold was determined by the lowest level at which an ABR can be recognized. ABR was recorded at 1-2 day before noise exposure and after noise exposure to assess the threshold shift.

Distortion product otoacoustic emission (DPOAE) was recorded by use of a Tucker-Davis R3 workstation with EC-1 high frequency speaker (Tucker-Davis Tech. Alachua, FL) (Zhu et al., 2013, 2015; Mei et al., 2017, Liu et al., 2023). Two pure tones (f_1_ and f_2_, f_2_/f_1_=1.22) were simultaneously delivered into the external ear canal through two plastic tubes and sealed with an earplug. The frequencies of two-testing sounds were determined by a geometric mean of f_1_ and f_2_ (f_0_ = (f_1_ × f_2_)^1/2^) at f_0_ = 4, 8, 16, and 20 kHz with f_2_/f_1_ = 1.22. The intensity of f_1_ (I_1_) was set at 5 dB SPL higher than that of f_2_ (I_2_). The distortion product was recorded with an average of 150 times and a cubic distortion product of 2f_1_ – f_2_ was measured as DPOAE.

### Cochlear microphonic (CM) recording

CM was evoked by 8-24 kHz tone bursts and recorded with the same electrode setting as ABR recording as previously reported (Zhu et al., 2013; Liu et al., 2023). The signal was amplified by 50,000 with 3-50 kHz filter and averaged by 100 times.

### Noise exposure

As described in our previous papers (Zhao et al., 2021, 2022; Liu et al., 2023), mice were awake in a small cage under loud-speakers in a sound-proof chamber and exposed to white-noise (95-98 dB SPL) for 2 hr, one time. Sound pressure level and spectrum in the cage were measured prior to placement of the animal.

### Patch-clamp recording and nonlinear capacitance measurement

OHCs were freshly isolated from the mouse cochlea (Zhu et al., 2013). The classical whole-cell recording was performed. A patch pipette was filled with an intracellular solution (140 KCl, 10 EGTA, 2 MgCl2, 10 HEPES in mM; 300 mOsm and pH 7.2) and had an initial resistance of 2.5–3.5 MΩ in the bath solution (130 NaCl, 5.37 KCl, 1.47 MgCl_2_, 2 CaCl_2_, and 10 HEPES in mM; 300 mOsm and pH 7.2) (Yu and Zhao, 2008). The pipette was patched at the basal nuclear pole of the OHC under a whole-cell configuration using Axopatch 200B patch clamp amplifier (Molecular Devices, CA) with jClamp (Scisft, New Haven, CT). Nonlinear capacitance (NLC) was measured by a two-sinusoidal method and fitted to the first derivative of a two-state Boltzmann function with jClamp and MATLAB (Yu and Zhao, 2008; Zhu et al., 2013).

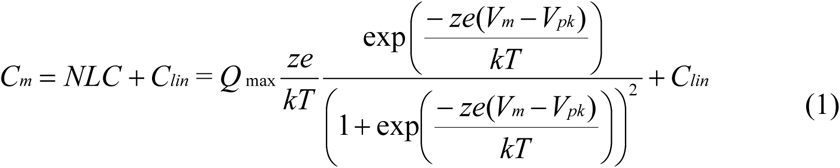

where *Q_max_* is the maximum charge transferred, *V_pk_* is the peak of NLC, *z* is the number of elementary charge (*e*), *k* is Boltzmann’s constant, and *T* is the absolute temperature. Membrane potential (*V_m_*) was corrected for electrode access resistance (*R_s_*).

### Reproducibility, data processing, and statistical analysis

The numbers of recording mice in each experiment were indicated in the related figure. Each experiment was repeated at least three times. Data were plotted by SigmaPlot and statistically analyzed by SPSS (SPSS Inc.; Chicago, IL). Error bars represent SEM. Parametric and nonparametric data comparisons were performed using one-way ANOVA or Student t tests after assessment of normality and variance (SPSS, SPSS Inc). The threshold for significance was α = 0.05. ANOVAs used Bonferroni *post hoc* test.

## Results

### P2x7 expression in the cochlea

We first examined P2x7 expression in the cochlea (Fig. 1). Immunofluorescent staining shows that P2x7 had high expression at SG neurons (Fig. 1A) and synapse areas under IHCs and OHCs with MOC and LOC fibers (Fig. 1B-D). In particularly, the P2x7 labeling mainly appeared at the modiolar side of IHCs (Fig. 1B). The P2x7 labeling located near the CtBP2-labeled ribbons under IHCs (Fig. 1E&F). However, the P2x7 labeling and CtBP2 labeling were not jointed together (Fig. 1F). We further used the *Peripherin-*eGFP mouse, in which type II SG neurons are labeled with eGFP, to examine P2x7 expression and found that type II SG neurons had intensive expression of P2x7 (Fig. 1I-K). The labeling of P2x7 is visible not only at the type II SG neuron soma (Fig. 1I) but also at its fibers (Fig. 1J&K). Finally, we examined P2x7 staining in P2x7 KO mice; no P2x7 labeling is visible in P2x7 KO mice (Fig. 1G&H). In comparison with WT mice, the MOC fibers in the P2x7 KO mice had no apparent alternations (Fig. 1G&H).

**Fig. 1.**
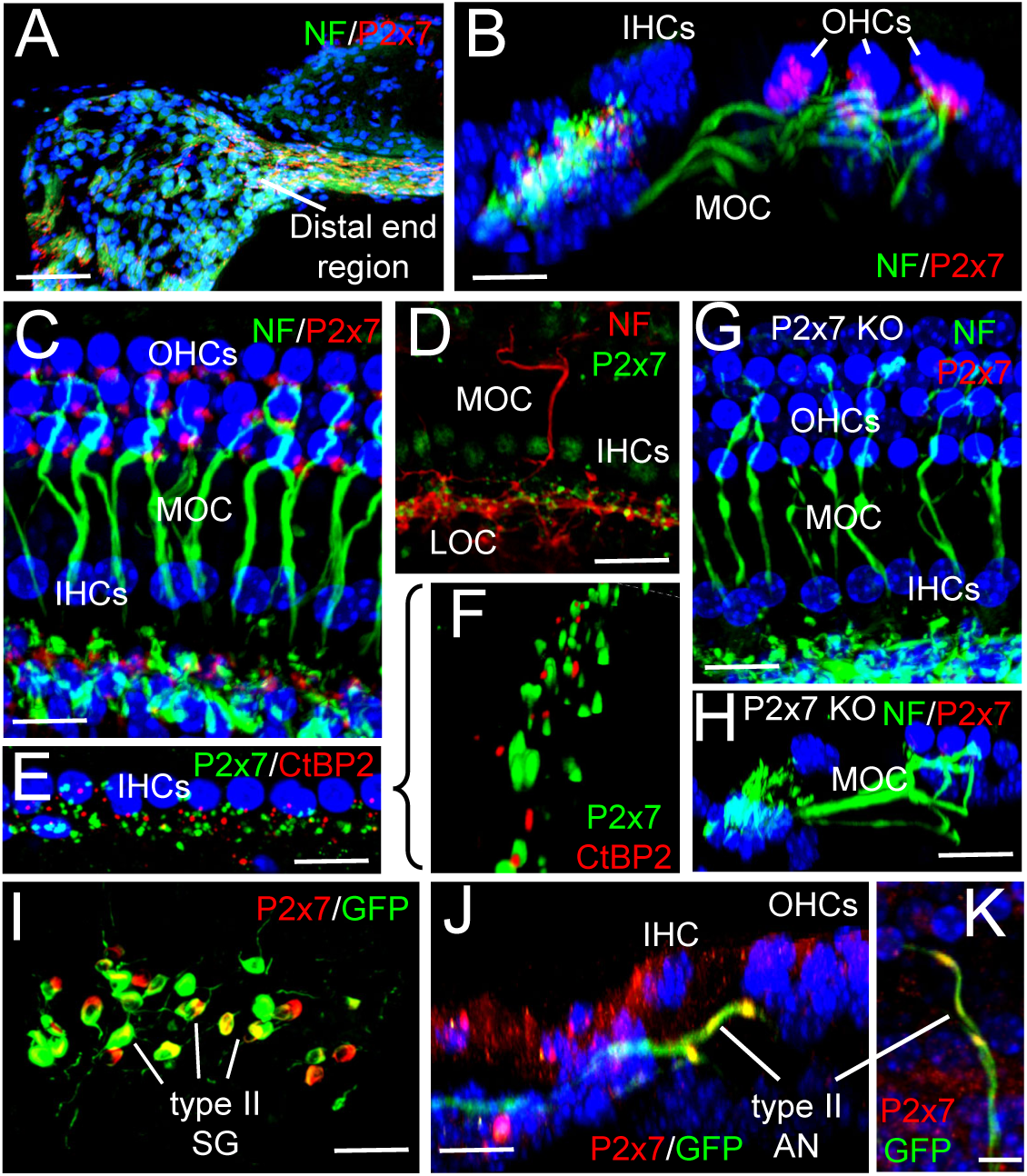
Immunofluorescent staining for P2x7 expression in the cochlea. **A&B**: P2x7 labeling in the cochlear cross-section or cross section view. The intensive staining of P2x7 is visible in the spiral ganglion (SG) neurons at the distal end region and the synapse areas under inner hair cells (IHCs) and outer hair cells (OHCs). **C&D**: P2x7 labeling in the whole-cochlear mounting preparation. P2x7 labeling is visible at MOC and LOC fibers. **E&F**: P2x7 labeling in the synapse area under IHCs with co-staining of CtBP2 for ribbon synapses. **G&H**: No P2x7 labeling in P2x7 KO mice. **I-K**: P2x7 expression in type II SG neurons with eGFP labeling in *Peripherin*-eGFP transgenic mice. Type II SG neurons and fibers have strong P2x7 labeling. Scale bar: 50, 30, 20, and 10 µm in penal A, B-G, H-J, and K, respectively.

### Enhanced acoustic responses in P2x7 KO mice

Deletion of P2x7 increased acoustic responses (Fig. 2). To assess acoustic responses, we first recorded acoustic startle response (ASR). The ASR in P2x7 KO mice was significantly increased (Fig. 2B&C). In P2x7 KO mice, the peaks of ASR at 80, 90, 100, and 110 dB SPL were 385.9±82.90, 666.7±142.7, 3371±589.5, and 6525±611.4 mV, respectively. In comparison with 276.6±58.76, 313.3±56.08, 1701±513.2, and 2230±231.8 mV in WT mice, the ASR in P2x7 KO mice was increased by 139.5±30.0, 212.8±45.6, 198.2±34.6, and 292.6±27.4% at 80, 90, 100, and 100 dB SPL, respectively. The increment was increased as the stimulus intensity increased. However, there was no significant difference in latency between P2x7 KO and WT mice (Fig. 2D).

**Fig. 2.**
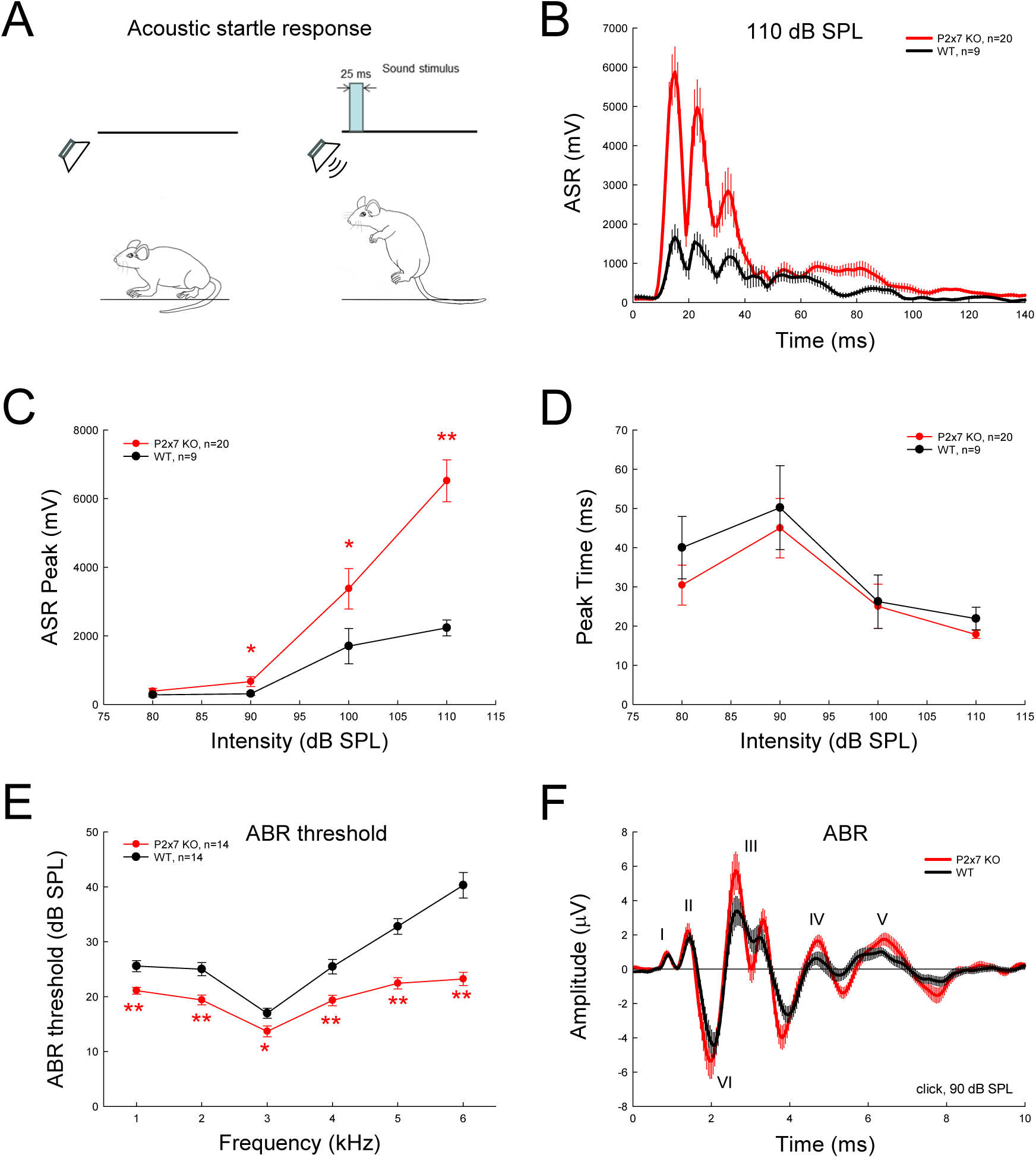
Enhanced acoustic responses in P2x7 KO mice. **A-D**: Enhanced acoustic startle response (ASR) in the P2x7 KO mice. Penal **A** is a diagram of ASR recording. The ASR in P2x7 KO mice is increased in comparison with that in wild-type (WT) mice (panel **B&C**). However, there is no significant difference in the latency of ASRs between P2x7 KO mice and WT mice (panel **D**). **E-F:** Enhanced auditory brainstem response (ABR) in P2x7 KO mice. Panel **F** is the average of ABR waveforms evoked by 90 dB SPL of click stimulations. I, II, III, IV, V, and VI represent the peak of ABR waveforms I, II, III, IV, V, and VI, respectively. *: P < 0.05, **: P < 0.01, t test, 2-tailed.

### Increase of auditory brainstem responses in P2x7 KO mice

ABR in P2x7 KO mice was also enhanced (Fig. 2E&F). The thresholds of ABR in P2x7 KO mice were significantly lower than those in WT mice (Fig. 2E). At the suprathreshold response, ABR waveforms in P2x7 KO mice appeared larger than that in WT (Fig. 2F). The quantitative measurements show that except peak I and V, ABR peaks II, III, IV, and VI evoked by 90 dB SPL clicks in P2x7 KO mice were significantly increased (Fig. 3). In particular, the amplitudes of peak II, III, and VI in P2x7 KO mice had ∼150% of increments in 60-110 dB SPL of tested intensity range (Fig. 3B, C, & F). However, there were no significant changes in peak latencies between P2x7 KO and WT mice (Fig. S1).

**Fig. 3.**
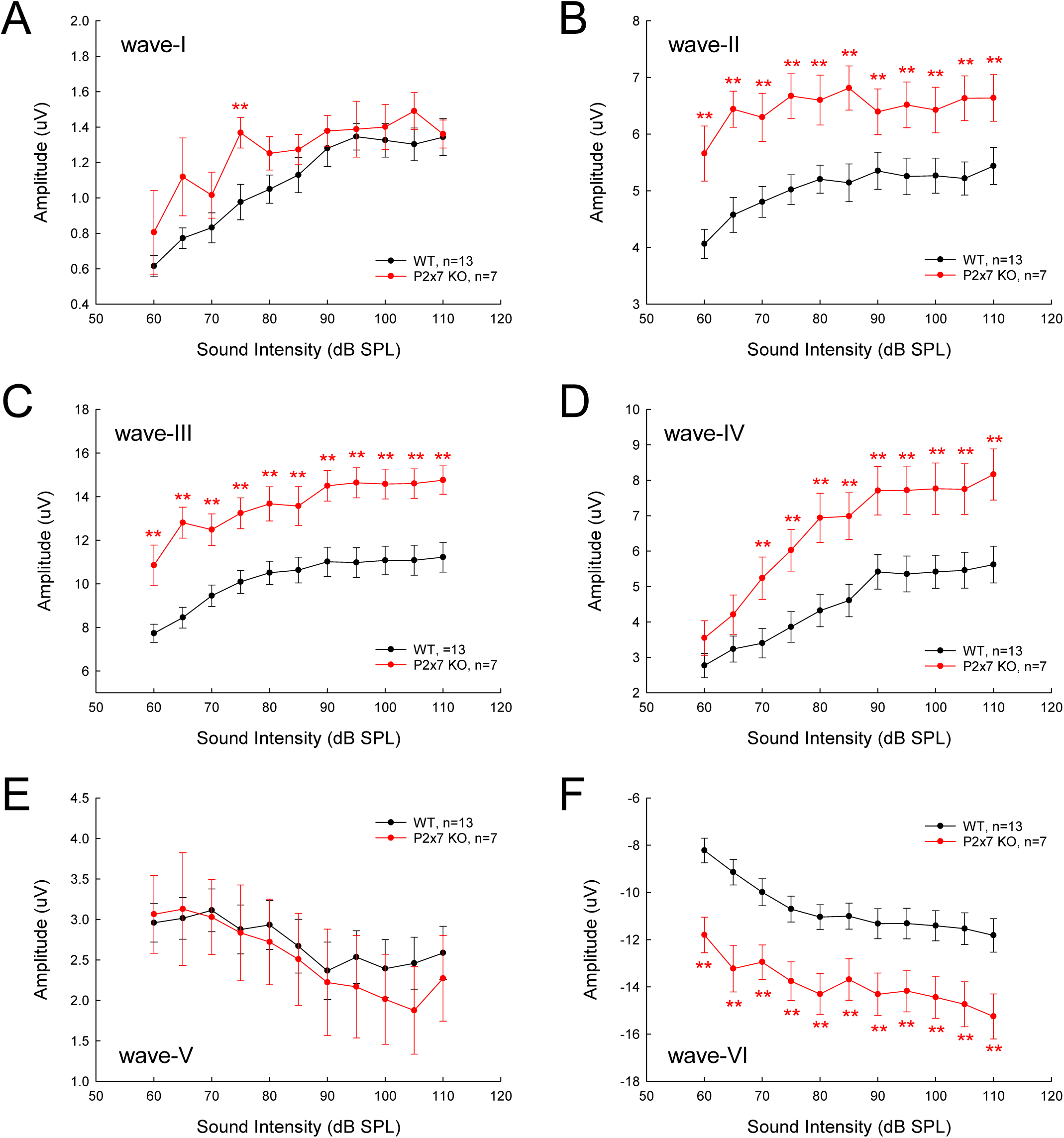
Quantitative measurement of ABR increment in P2x7 KO mice. ABRs were evoked by 90 dB SPL click stimuli. The amplitudes of wave II-VI in P2x7 KO mice significantly increased in comparison with those in WT mice. *: P<0.05, **: P<0.01, t test, 2-tailed.

### Increase of cochlear microphonics in P2x7 KO mice

We further examined the auditory receptor current and found that cochlear microphonics (CM) in P2x7 KO mice was also increased (Fig. 4). CM increases were visible in all tested intensity (60-100 dB SPL) and tested frequency ranges (8-24 kHz) (Fig. 4B-D). In comparison with WT mice, the increases of CM in P2x7 KO mice were increased as frequency increased (Fig. 4E). The increase was also increased as the intensity increased (Fig. 4E). At 24 kHz, CM in P2x7 KO mice at 100 dB SPL was increased by ∼500% (Fig. 4E&F).

**Fig. 4.**
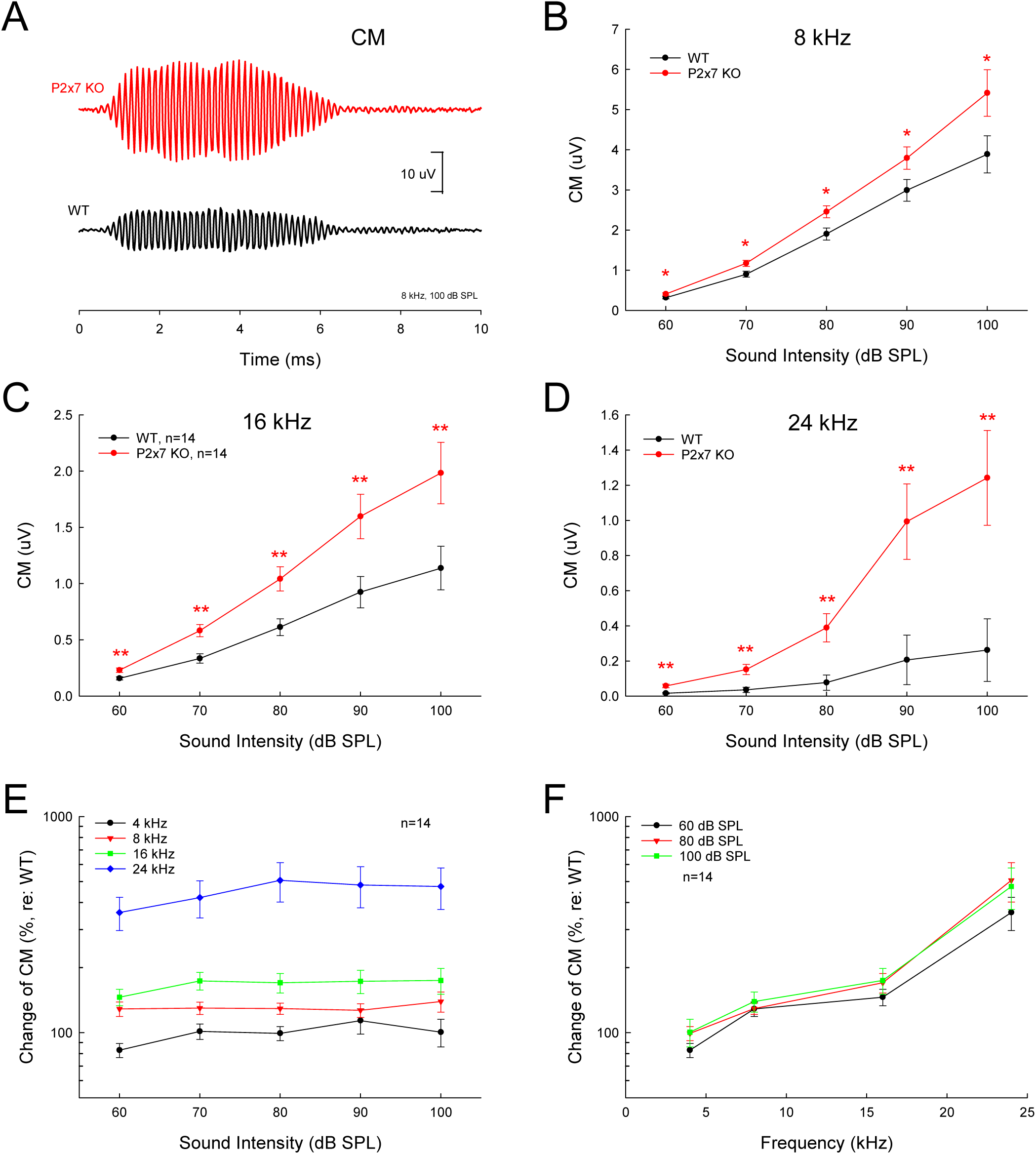
Increase of CM in P2x7 KO mice. **A**: CM traces recorded from P2x7 KO and WT mice. CM was evoked by 100 dB SPL, 8 kHz tone bursts. **B-D**: CMs in P2x7 KO mice were significantly increased in comparison with those in WT mice in the tested 8-24 kHz frequency range. Mice were 2-3 months old. **E:** Input-output (I-O) function of CM in P2x7 KO mice. The CMs in P2x7 KO mice were normalized to those in WT mice. The increase of CM in P2x7 KO mice retains constant in 60-100 dB SPL of the tested intensity range. **F**: The CM in P2x7 KO mice increases as the frequency increasing. *: P<0.05, **: P<0.01, t test, 2-tailed.

### Changes of outer hair cell electromotility in P2x7 KO mice

MOC fibers target to OHCs and P2x7 had intensive expression at the OHC synaptic ending (Fig. 1B). We further examine the effect of P2x7 KO on OHC electromotility. OHC electromotility associated nonlinear capacitance (NLC) was recorded. Fig. 5 shows that after deletion of P2x7, NLC was increased and shifted to the negative hyperpolarization direction. In comparison with voltage of peak capacitance (V_pk_) in WT mice (−41.1±3.26 mV), the V_pk_ in P2x7 KO mice significantly shifted to −66.2±6.31 mV (Fig. 5B); Q_max_ and z were changed from 0.39±0.10 pC and 0.94±0.07 in WT mice to 0.86±0.09 pC and 0.66±0.04 in P2x7 KO mice (Fig. 5C&D), i.e., NLC was increased from 3.44±0.74 pF in WT mice to 5.18±0.31 pF in P2x7 KO mice. However, linear capacitance (C_lin_), which is mainly determined by cell length, in WT and P2x7 KO mice was 5.82±0.31 and 5.82±0.51 pF, respectively and had no significant change.

**Fig. 5.**
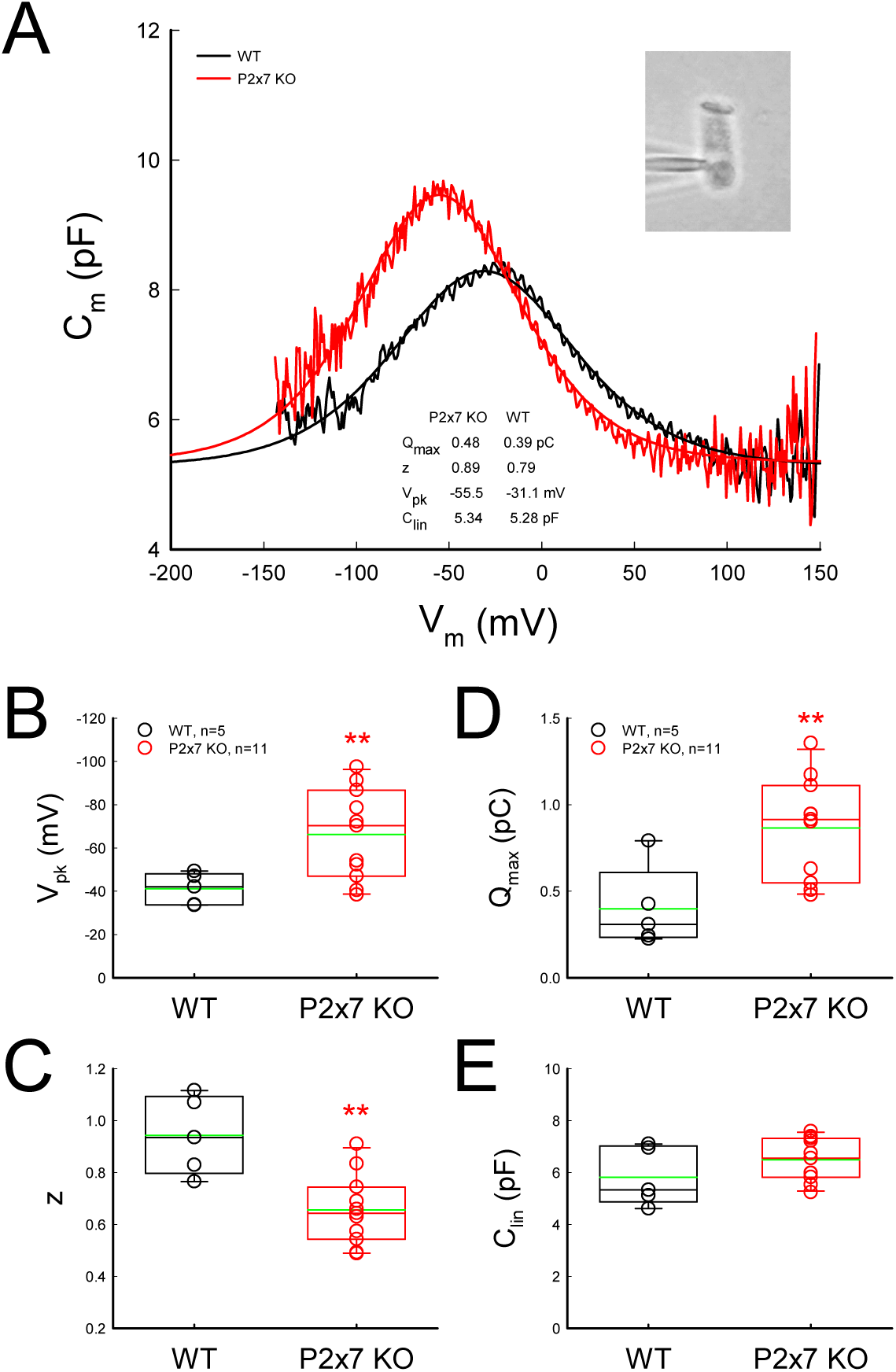
Increase of outer hair cell electromotility associated nonlinear capacitance (NLC) in P2x7 KO mice. **A**: NLC recorded from P2x7 KO and WT mice. Smooth lines represent fitting by the first derivative of Boltzmann equation. The parameters of fitting are Q_max_=0.48 & 0.39 pC; z=0.89 & 0.79; V_pk_=-55.5 & −s31.1 mV; and C_lin_=5.34 & 5.28 pF for P2x7 KO and WT mice, respectively. Inset: A captured image of patch clamp recording in a mouse OHC. **B-E**: Parameters of NLC fitting in P2x7 KO and WT mice. Green lines in boxes represent the meaning level. Mice were P30-45 old. In P2x7 KO mice, V_pk_ was shifted to negative potential, Q_max_ was increased, and z was reduced. However, there was no significant difference in C_lin_, which is mainly determined by cell length. *: P < 0.05, **: P < 0.01, t test, 2-tailed.

### Increase of susceptibility to noise in P2x7 KO mice

The cochlear efferent system provides negative feedback to control hearing sensitivity and protect from noise trauma by reducing responses to acoustic stimulation. We further examined the effect of P2x7 deficiency on the protection from noise trauma. Fig. 6 shows that after exposure to 98-100 dB SPL of white noise for 2 hours, ABR thresholds in P2x7 KO mice had significant increase in comparison with those in WT mice. At post-exposure day 1 (P1), ABR thresholds of P2x7 KO mice were elevated to 60-90 dB SPL in comparison with 40-60 dB SPL in WT mice (Fig. 6C). In comparison with those in pre-exposure, the increases of ABR thresholds in P2x7 KO and WT mice were 30-50 dB and 10-30 dB, respectively (Fig. 6D). The increase in middle-high frequency range also appeared large (Fig. 6C&D). Then, the ABR thresholds in WT mice were quickly recovered (Fig. 6A&B). At P3, the thresholds in WT mice already returned to the pre-exposure levels, whereas the ABR thresholds in P2x7 KO mice only showed a small, slowly recovering (Fig. 6A&B). At P28, the ABR thresholds in P2x7 KO mice still retained above 40-80 dB SPL (Fig. 6E) and had 15-40 dB of threshold shift (Fig. 6F), i.e., 15-40 dB permanent threshold shift (PTS). The shifting was also large at middle-high frequency range (Fig. 6F).

**Fig. 6.**
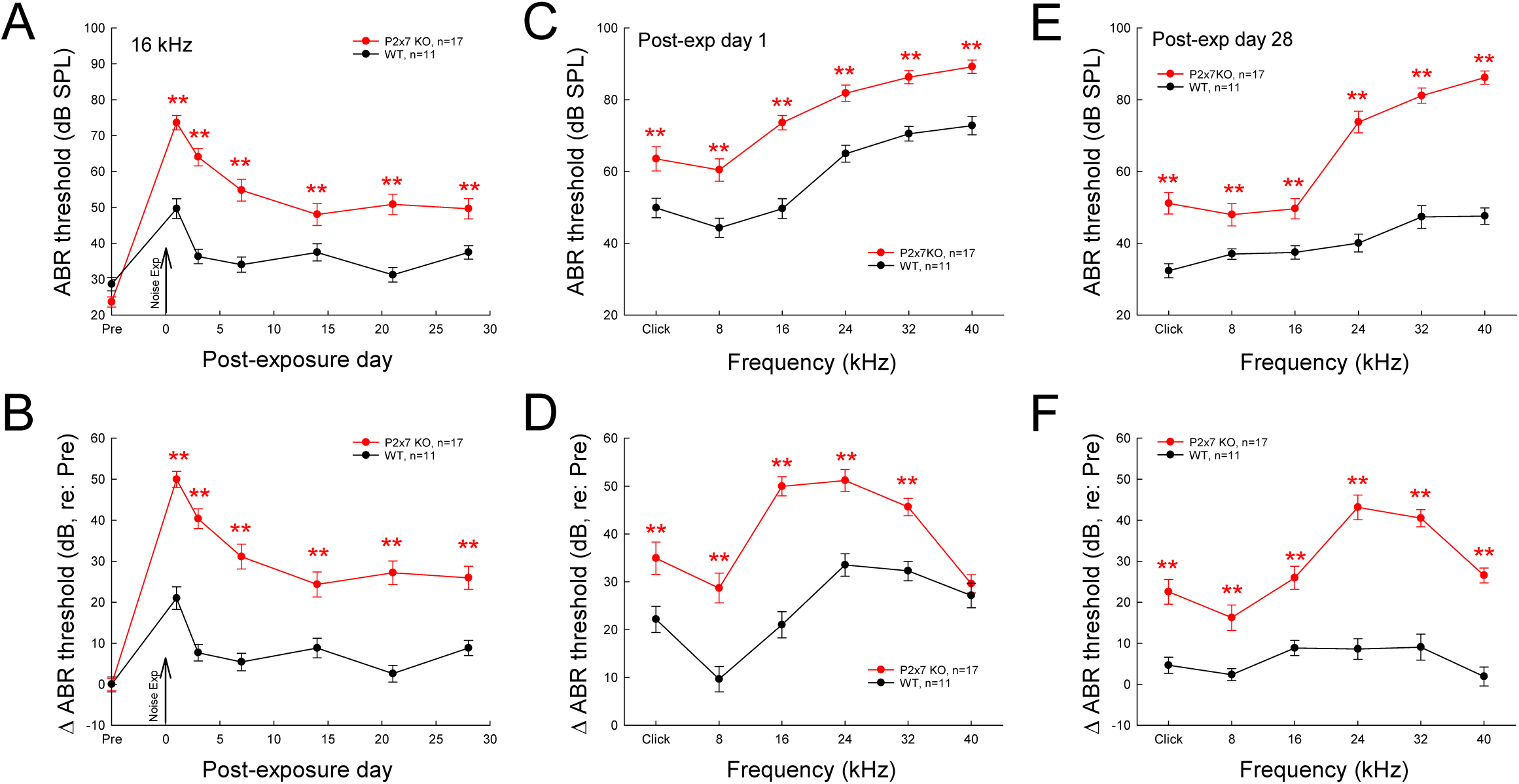
Knockout of P2x7 increases susceptibility to noise trauma. Both P2x7 KO and WT mice were exposed to 98-100 dB SPL white noise for 2 hrs, one time. The noise-expose day is defined as day 0 (P0). **A-B**: Dynamic changes of ABR thresholds in P2x7 KO and WT mice after noise exposure. Vertical arrows indicate the noise exposure day (P0). ABR were evoked by 16 kHz tone bursts. ABR thresholds in panel **B** were normalized to the pre-noise exposure levels to reveal changes in the ABR threshold after noise exposure. In comparison with WT mice, ABR thresholds in P2x7 KO mice were not completely recovered after noise exposure. **C-D**: Changes of ABR thresholds in frequency range in P2x7 KO and WT at post-exposure day 1 (P1). In comparison with WT mice, P2x7 KO mice have large ABR threshold increased. **E-F**: Increases of ABR thresholds in frequency range in P2x7 KO at P28. In comparison with WT mice, the ABR thresholds in P2x7 KO mice have ∼ 20-40 dB SPL permanent threshold shift (PTS); the PTS appears large at high frequency range. *: P < 0.05, **: P < 0.01, t test, 2-tailed.

### Noise exposure impairing acoustic emission in P2x7 KO mice

Noise also significantly impaired acoustic emissions in P2x7 KO mice (Fig. 7). Before noise exposure, distortion product of acoustic emission (DPOAE) in P2x7 KO mice was no significant difference from WT mice (Fig. 7 C&D). After noise exposure, DPOAEs in P2x7 KO mice at middle and high frequency range almost disappeared and did not recover (Fig. 7E-H), while WT mice only had slightly reduction at P1 (Fig. 7G&H) and quickly recovered to the pre-exposure levels at P3 (Fig. 7E-H).

**Fig. 7.**
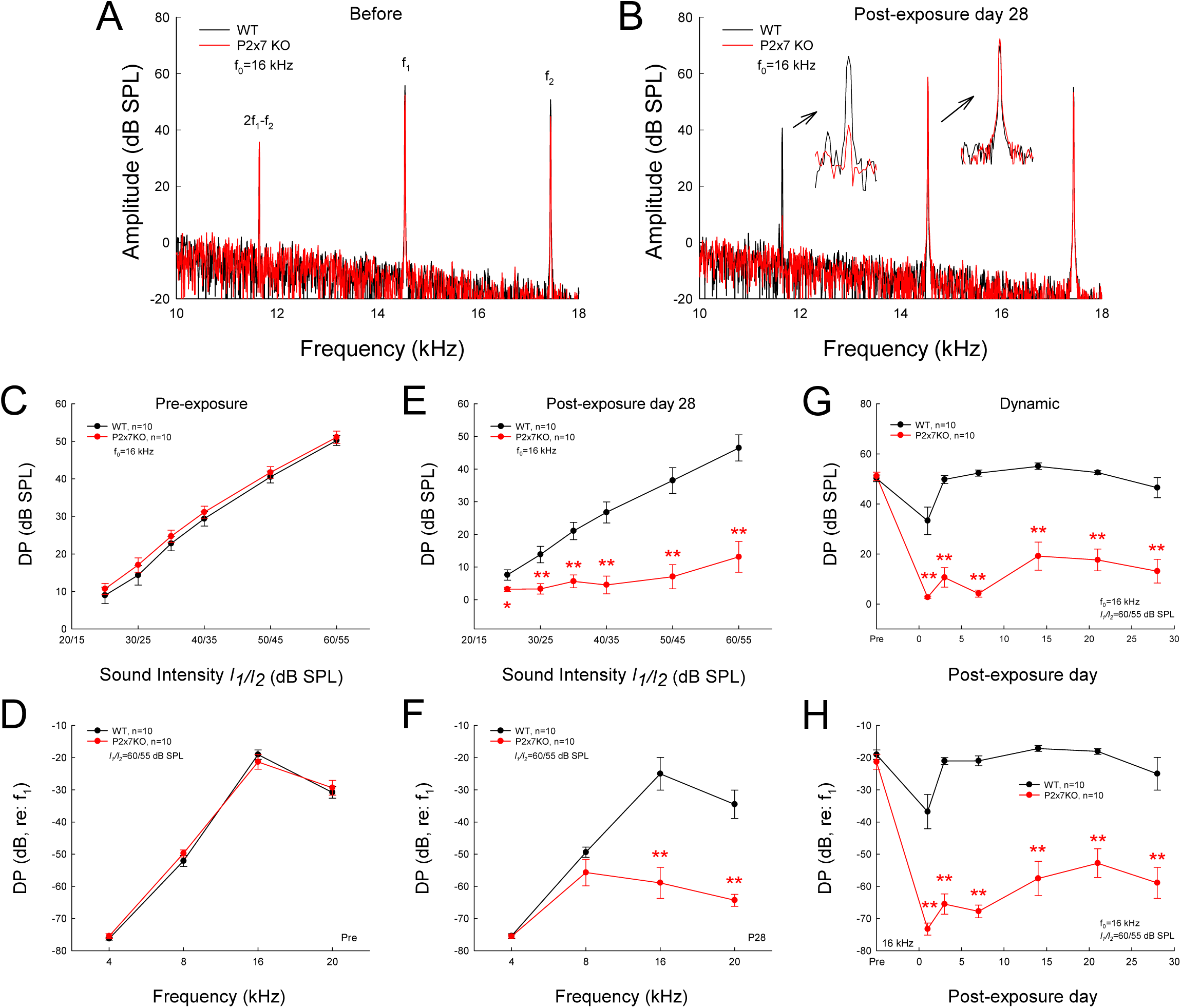
Reduction of DPOAE in P2x7 KO mice after noise exposure. **A-B**: Spectra of acoustic emission recorded from WT and P2x7 KO mice before noise exposure and at post-exposure day 28 (P28). DPOAEs were evoked by two-tone stimulation. f_0_=16 kHz, I_1_/I_2_=60/55 dB SPL. Insets: Large scale plots of 2f_1_-f_2_ and f_1_ peaks at P28. **C-D**: DPOAE of P2x7 KO and WT mice before noise exposure. In panel **C**, the DPOAE is normalized to the peak of f_1_. There is no significant difference between P2x7 KO and WT mouse groups. **E-F**: Reduction of DPOAE in P2x7 KO mice after noise exposure. DPOAEs were recorded at P28. In comparison with WT mice, DPOAEs at high frequency in P2x7 KO mice were significantly reduced after noise exposure. **G-H**: Dynamic changes of DPOAE in P2x7 KO and WT mice after noise exposure. DPOAEs in P2x7 KO mice are not recovered and have ∼ 40 dB reduction in comparison with WT mice after noise exposure. *: P < 0.05, **: P < 0.01, t test, 2-tailed.

We further examined changes of ribbon synapses in IHCs for noise exposure (Fig. 8). In the control groups without noise exposure, the ribbon labeling puncta per IHC in WT and P2x7 KO mice were 10.5±0.93 and 10.9±0.97/IHC, respectively (Fig. 8C). There was no significant difference between WT and P2x7 KO mice (P=0.74, 2-tailed t test). However, after noise exposure, the ribbon puncta in WT and P2x7 KO mice were significantly reduced to 6.40±0.78 and 3.77±0.50/IHC (P <0.05, 2-tailed t test), respectively. The reduction of ribbons in P2x7 KO mice was almost twice more than that in WT mice (P=0.03, 2-tailed t test).

**Fig. 8.**
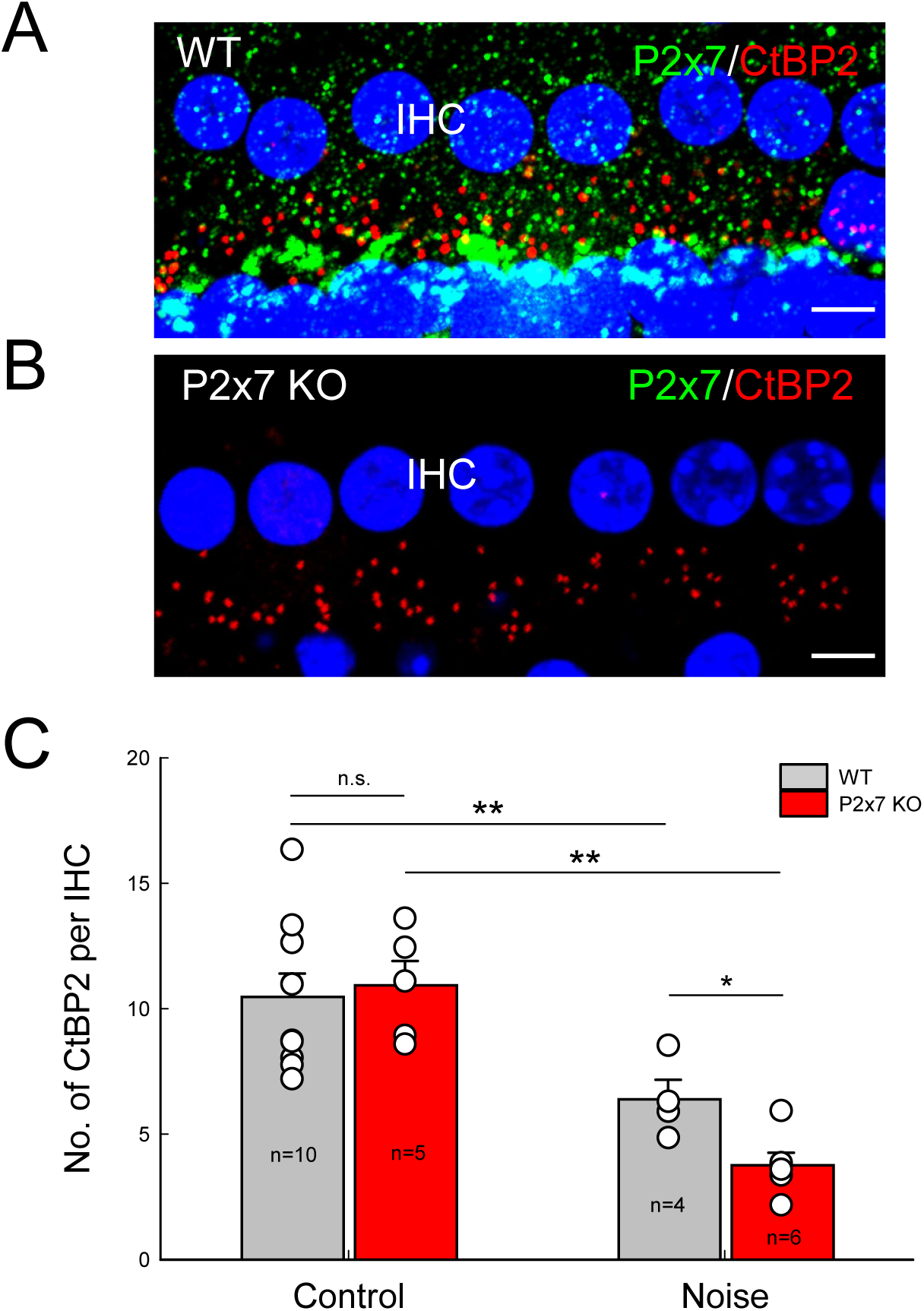
More significant reduction of ribbon synapse in P2x7 KO mice after noise exposure. **A-B:** Immunofluorescent staining for P2x7 and CtBP2 in WT and P2x7 KO mice. Scale bar: 10 µm. **C:** Quantitative accounts of CtBP2 puncta per IHC. In control groups without noise exposure, there is no significant difference of CtBP2 labeling puncta at the IHC between WT and P2x7 KO mice. After noise exposure at post-exposure day 28 (P28), CtBP2 labeling puncta in both WT and P2x7 KO mice had significant reduction in comparison with control groups. The P2x7 KO mice appear more significant reduction than that in WT mice. *: P < 0.05, **: P < 0.01, t test, 2-tailed.

## Discussion

In this study, we found that P2x7 had extensively expressions at type II ANs and MOC and LOC nerves in the cochlea (Fig. 1). Deletion of P2x7 led to hearing over-sensitivity (Figs. 2-4) and susceptibility to noise (Figs. 6-8) by enhanced OHC electromotility (Fig. 5). These findings suggest that P2x7 receptors have a critical role in type II AN function and is required for the cochlear efferent system controlling hearing sensitivity and protection from noise trauma.

It is critical for hearing that the cochlear efferent system provides negative feedback to hair cells to regulate hearing sensitivity and protect against acoustic trauma. The MOC nerves project to OHCs, whereas the LOC nerves project to IHCs and form synapses with the dendrites of type I afferent auditory nerves under the IHCs. The MOC fibers are cholinergic fibers and acetylcholine (ACh) is a primary neurotransmitter, and the LOC has cholinergic fibers and dopaminergic fibers, releasing ACh, dopamine, and other neurotransmitters (Eybalin, 1993; Lustig, 2006; Maison et al., 2012). In this study, we found that P2x7 has extensive expression at the synapse areas under IHCs and OHCs with LOC and MOC fibers (Fig. 1), suggesting that P2x7-mediated ATP purinergic signaling pathway also play a critical role in the cochlear efferent system to mediate hair cell activities.

Indeed, we found that deletion of P2x7 led to ASR increased, ABR threshold reduced, and the suprathreshold ABR responses increased (Figs. 2-3); CM was also increased (Fig. 4). Such hearing over-sensitive responses were consistent with the consequent results from the impairment and deficiency of the cochlear efferent negative feedback induced by P2x7 KO. We previously reported that MOC fibers have innervations on gap junctions between the cochlear supporting cells to mediate OHC function, which is especially important for “slow” efferent inhibition (Zhao et al, 2022); deficiency of Cx26 gap junctions also could induce hearing over-sensitivity by enhancing OHC electromotility (Liu et al., 2023). Thus, the deficiency of the cochlear efferent system can lead to hearing over-sensitivity by enhancing active cochlear mechanics.

P2x7 receptors had intensive labeling in the OHC synapse area (Fig. 1). We found that deletion of P2x7 increased OHC electromotility-associated NLC and shifted it to hyperpolarization (Fig. 5). Currently, the detailed mechanism underlying control of OHC electromotility is largely undetermined. We previously reported that ATP can mediate OHC electromotility through activation of P2x receptors to influx Ca^++^ (Yu and Zhao, 2008). ATP can activate P2x7 receptors to influx K^+^ and Ca^++^ ions inhibiting cell depolarization (Surprenant and North, 2009). Gap junction hemichannels in the cochlear supporting cells can release ATP to control hearing sensitivity (Zhao et al., 2005, Chen et al., 2015); the release increases as the mechanical stimulation (sound intensity) increases, which in turn mediates OHC electromotility to reduce the gain of the cochlear amplifier (Zhao et al., 2005). Deletion of P2x7 did not have significant changes in DPOAE under normal condition (Fig. 7C&D). However, P2x7 KO mice was vulnerable to noise exposure inducing active cochlear mechanics impaired (Fig. 7E-H). Deficiency of P2x7 also had no hearing loss (Fig. 2E) but increased susceptibility to noise trauma resulting in permanent hearing loss (Fig. 6). These data demonstrated that deletion of P2x7 could impair the cochlear efferent function and then impair the gain control of active cochlear amplification and OHC electromotility leading to susceptibility to noise and noise-induced hearing loss.

Type II ANs innervate OHCs. Since it is difficult to be identified and recorded, little is known about channel information in the type II SG neurons and nerves. In this study, we found that P2x7 receptors have intensive labeling in type II SG neurons and fibers (Fig. 1), suggesting that P2x7 receptors have a critical role in the function of type II ANs. This is also consistent with previous reports that ATP can stimulate type II AN activation (Weisz et al., 2009) to sense auditory nociception (Flores et al., 2015; Liu et al., 2015). However, type II AN function should not be limited to nociception sensing, since type II ANs can also be activated by nondamaging sounds as well (Weisz et al., 2014, 2021). Indeed, the type II AN is also considered to provide an important input for the cochlear efferent system negative loop to control active cochlear amplifier OHC electromotility (Froud et al., 2015), although it had a debate (Maison et al., 2016). Our results support this concept. We found that P2x7 has intensive labeling in type II auditory nerves (Fig. 1); deletion of P2x7 impaired the efferent negative control to increase hearing sensitivity (Figs. 2-4) and lead to susceptibility to noise (Figs. 6-8), suggesting that P2x7 is not only required for the cochlear efferent function and is also critical for providing input information to the efferent system to control OHC electromotility.

## Acknowledgements

This work was supported by NIH R01 DC017025, DC019687, and RF1 AG082216 to HBZ.

## Author Contributions

HBZ conceived the general framework of this study. CL, SF, TYZ, JC, LML, YN, and HBZ performed the experiments, analyzed data, and wrote the paper. All authors reviewed the manuscript and provided the input.

## Supplementary figures

**Fig. S1.**
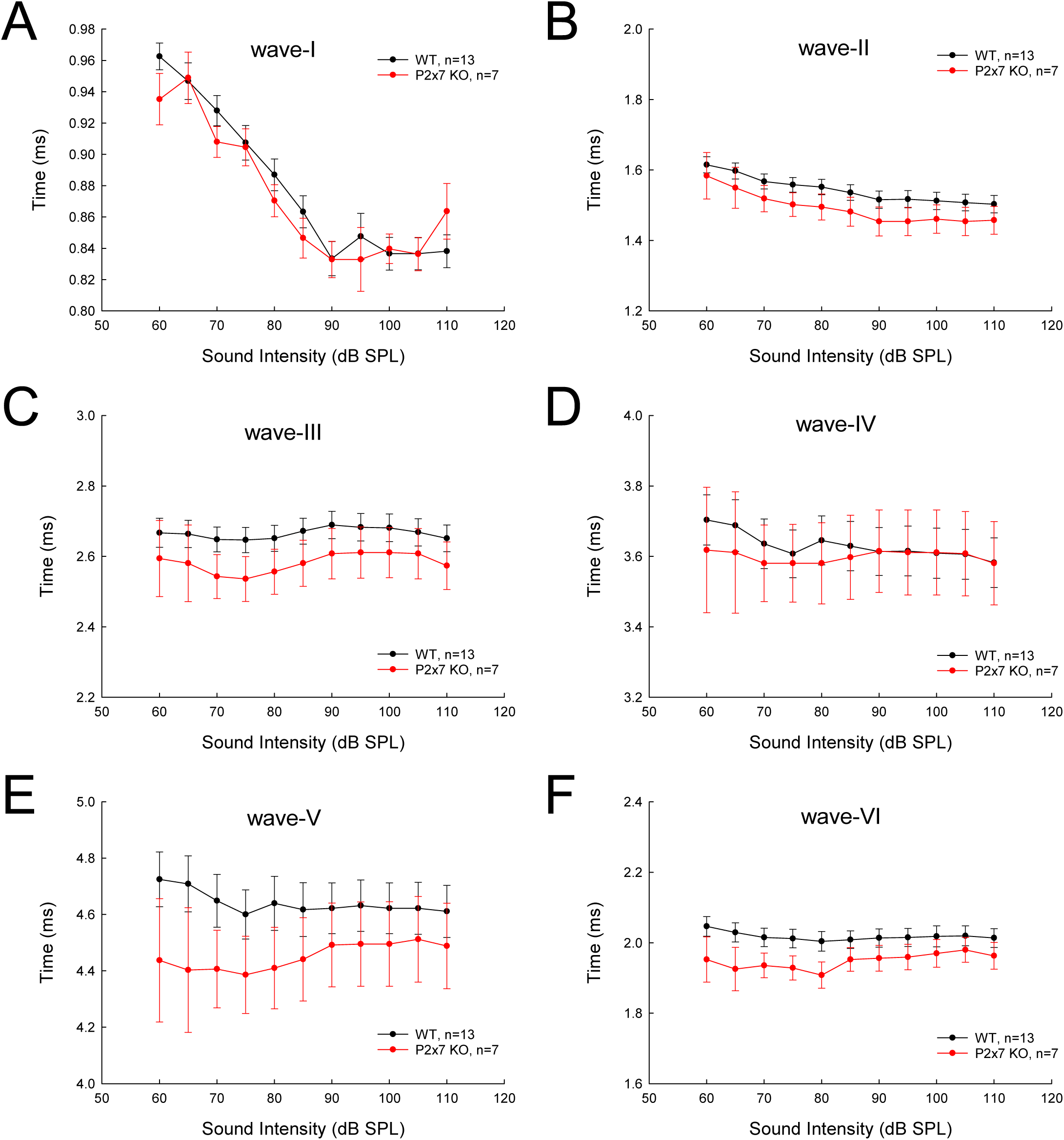
No significant changes of the latency of ABR in P2x7 KO mice. Mice were 2-3 months old. ABRs were evoked by 90 dB SPL of click stimuli. The peak times of ABR waveform I, II, III, IV, V, and VI were measured and averaged (n= 7 and 13 for P2x7 KO and WT mice, respectively). The ABR peak times in P2x7 KO mice appear smaller than those in WT mice. However, there are no significant differences between WT and P2x7 KO mice (P > 0.05, t test).

